# The E3 ligase Thin controls homeostatic plasticity through neurotransmitter release repression

**DOI:** 10.1101/2021.06.16.448554

**Authors:** Martin Baccino-Calace, Katharina Schmidt, Martin Müller

## Abstract

Synaptic proteins and synaptic transmission are under homeostatic control, but the relationship between these two processes remains enigmatic. Here, we systematically investigated the role of E3 ligases, key regulators of protein degradation-mediated proteostasis, in presynaptic homeostatic plasticity (PHP). An electrophysiology-based genetic screen of 157 E3 ligase-encoding genes at the *Drosophila* neuromuscular junction identified *thin*, an ortholog of human *tripartite motif-containing 32* (*TRIM32*), a gene implicated in several neural disorders, including Autism Spectrum Disorder and schizophrenia. We demonstrate that *thin* functions presynaptically during rapid and sustained PHP. Presynaptic *thin* negatively regulates neurotransmitter release under baseline conditions by limiting the number of release-ready vesicles, independent of gross morphological defects. We provide genetic evidence that *thin* controls release through *dysbindin*, a schizophrenia-susceptibility gene required for PHP. Thin and Dysbindin localize in close proximity within presynaptic boutons, and Thin degrades Dysbindin *in vitro*. Thus, the E3 ligase Thin links protein degradation-dependent proteostasis of Dysbindin to homeostatic regulation of neurotransmitter release.

## INTRODUCTION

Nervous system function is remarkably robust despite continuous turnover of the proteins determining neural function. Work in nervous systems of various species has established that evolutionarily conserved homeostatic singling systems maintain neural activity within adaptive ranges ^1–3^. Chemical synapses evolved mechanisms that compensate for neural activity perturbations through homeostatic regulation of neurotransmitter release (‘presynaptic homeostatic plasticity’, PHP) ^2,4,5^, or neurotransmitter receptors (synaptic scaling) ^6^. Several studies have established links between homeostatic control of synaptic transmission and neural disease, such as Autism Spectrum Disorder ^7^, schizophrenia ^8^, or Amyotrophic Lateral Sclerosis ^9,10^. Synaptic proteins are continuously synthesized and degraded, resulting in half-lives ranging from hours to weeks ^11^. The Ubiquitin-Proteasome System (UPS) is a major protein degradation pathway that controls protein homeostasis, or proteostasis. E3 ubiquitin ligases confer specificity to the UPS by catalyzing the ubiquitination of specific target proteins, thereby regulating their function or targeting them for proteasomal degradation ^12^. Synaptic proteostasis, and E3 ligases in particular, have been implicated in various neural disorders ^13^. However, our understanding of the role of E3 ligases in the regulation of synaptic transmission is very limited. While several E3 ligases have been linked to postsynaptic forms of synaptic plasticity ^14^, only three E3 ligases, Scrapper ^15^, highwire ^16^ and Ariadne-1 ^17^ have been implicated in the regulation of presynaptic function. Moreover, a systematic investigation of E3 ligase function in the context of synaptic transmission is lacking.

PHP stabilizes synaptic efficacy in response to neurotransmitter receptor perturbation at neuromuscular junctions (NMJs) of *Drosophila melanogaster* ^2,4,5^, mice ^18^, rats ^19^, and humans ^20^. Furthermore, there is recent evidence for PHP in the mouse cerebellum ^21^. The molecular mechanisms underlying PHP are best understood at the *Drosophila* NMJ ^2^, because this system is amenable to electrophysiology-based genetic screens ^2,22,23^. At this synapse, pharmacological or genetic impairment of glutamate receptor activity triggers a retrograde signal that enhances presynaptic release, thereby precisely compensating for this perturbation ^4,5^. PHP can be induced within minutes after pharmacological receptor impairment ^5^. Severing the motoneuron axons forming the *Drosophila* NMJ in close vicinity of the synapse does not impair PHP ^5^, indicating that the mechanisms underlying PHP act locally at the synapse. Moreover, pharmacological inhibition of protein synthesis by cyclohexamide does not affect PHP at the *Drosophila* NMJ ^5^, suggesting that *de novo* protein synthesis is not required for PHP. By contrast, acute or sustained disruption of the presynaptic proteasome blocks PHP ^24^, demonstrating that presynaptic UPS-mediated proteostasis is required for PHP. Furthermore, genetic data link UPS-mediated degradation of two proteins, Dysbindin and RIM, to PHP ^24^. Yet, it is currently unclear how the UPS controls PHP. Based on the critical role of E3 ligases in UPS function, we hypothesized an involvement of E3 ligases in PHP.

Here, we realized an electrophysiology-based genetic screen to systematically analyse the role of E3 ligases in neurotransmitter release regulation and PHP at the *Drosophila* NMJ. This screen discovered that the E3 ligase-encoding gene *thin*, an ortholog of human *TRIM32* ^25,26^, controls neurotransmitter release and PHP. We provide evidence that *thin* regulates the number of release-ready synaptic vesicles through *dysbindin*, a gene linked to PHP in *Drosophila* and schizophrenia in humans.

## RESULTS

### An electrophysiology-based genetic screen identifies *thin*

To systematically test the role of E3 ligases in PHP, we first generated a list of genes predicted to encode E3 ligases in *D. melanogaster* (Figure 1A). To this end, we browsed the *D. melanogaster* genome for known E3-ligase domains ^27,28^. Moreover, we included homologs of predicted vertebrate E3-ligases (see Figure S1). This approach yielded 281 putative E3 ligase-encoding genes (Figure 1A), significantly higher than previously predicted for *D. melanogaster* (207 genes, ^27^). To explore the relationship between the number of E3 ligase-encoding genes and genome size, we plotted the number of putative E3 ligase-encoding genes over the total protein-coding gene number of three species, and compared it to the relationship between protein kinase-encoding genes and genome size (Figure 1A). The relatively constant ratio between the predicted number of E3 ligase-encoding genes and genome size across species (~0.02-0.03; Figure 1A; ^28^), suggests an evolutionarily conserved stoichiometry between E3 ligases and target proteins, similar to protein kinases (Figure 1A). Hence, a core mechanism of the UPS – protein ubiquitination – is likely conserved in *D. melanogaster*.

**Figure 1.**
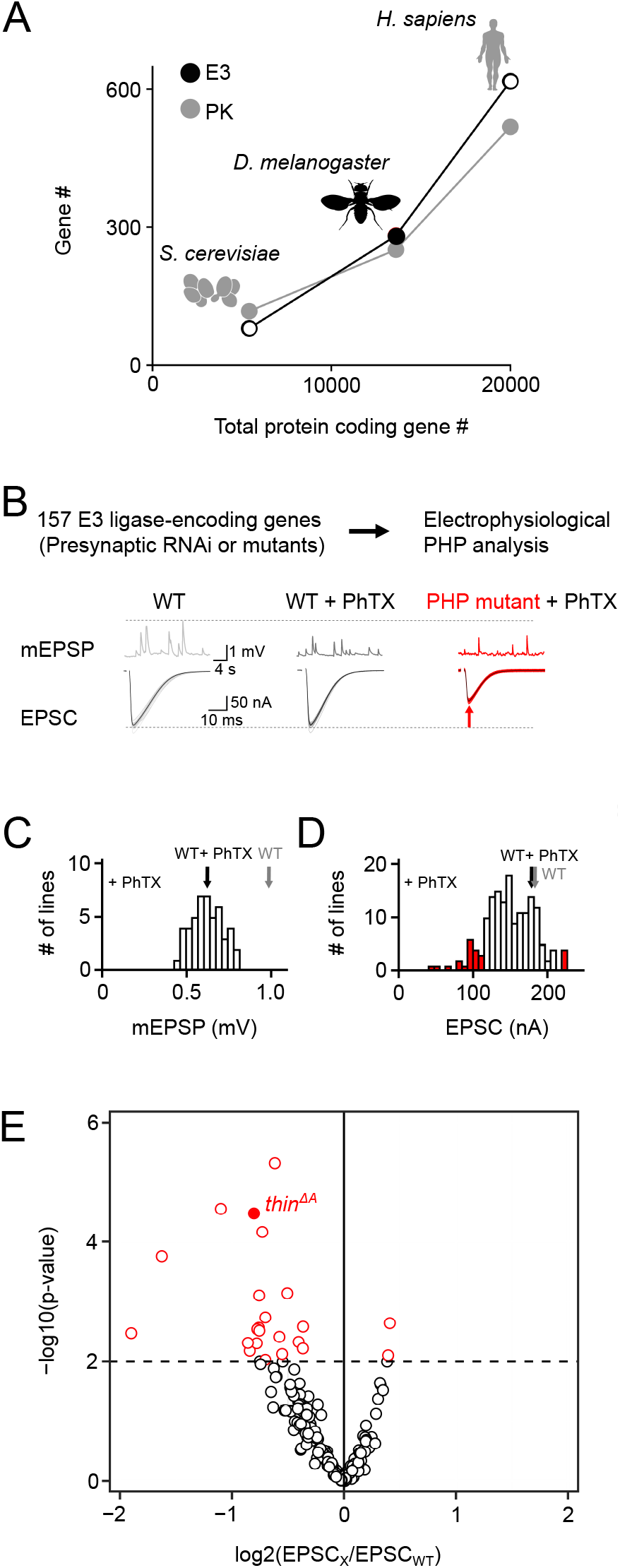
An electrophysiology-based genetic screen identifies *thin* as a synaptic homeostasis mutant. **A)** Number of putative E3 ligase-encoding genes (E3) and protein kinase-encoding genes (PK) as a function of total protein-coding gene number of *C. cerevisiae*, *D. melanogaster*, and *H. sapiens*. Note the similar relationship between E3 number, PK number and total protein-coding gene number across species. **B)** *Top:* 157 E3 ligase encoding genes and 11 associated genes (180 lines; presynaptic RNAi expression, *elav*^*c155*^ > *UAS-RNA*_*i*_, or mutants) were tested using two-electrode voltage clamp analysis at the *Drosophila* NMJ in the presence of the glutamate receptor antagonist PhTX-433 (‘PhTX’) to assess PHP (see Methods). *Bottom:* Exemplary mEPSPs and AP-evoked EPSCs recorded from WT controls, WT in the presence of PhTX (‘WT + PhTX’), and a PHP mutant in the presence of PhTX (‘PHP mutant + PhTX’). Note the decrease in mEPSP amplitude after PhTX treatment, indicating GluR inhibition, and the similar EPSC amplitude between WT and WT + PhTX, suggesting PHP. Small EPSC amplitudes in the presence of PhTX imply a defect in PHP or baseline synaptic transmission. **C)** Histogram of mean mEPSP amplitudes for each transgenic or mutant line (mean n=4, range 3-12; N=180 lines) following PhTX treatment. The wild-type (WT) averages under control conditions (‘WT’ n=16) and in the presence of PhTX (‘WT + PhTX’, n=16) are shown as gray and black arrows, respectively. **D)** Histogram of mean EPSC amplitudes (as in *C*). The red bars indicate transgenic or mutant lines with EPSC amplitudes significantly different to the WT control in the presence of PhTX. **E)** Volcano plot of the ratio between the mean EPSC amplitude of a transgenic or mutant line and WT (‘EPSC_x_/EPSC_WT_’) in the presence of PhTX (p values from one-way ANOVA with Tukey’s multiple comparisons). Transgenic or mutant lines with mean EPSC amplitude changes with p≤0.01 (dashed line) are shown in red. A deletion in the gene *thin* (*CG15105*; *thin*^*ΔA*^; LaBeau-DiMenna et al., 2012) that was selected for further analysis is shown as a filled red circle. One-way ANOVA with Tukey’s multiple comparisons was performed for statistical testing (C, D, E).

After prioritizing for evolutionarily-conserved genes that were shown or predicted to be expressed in the nervous system (Figure S1), we investigated PHP after genetic perturbation of 157 putative E3 ligase genes and 11 associated genes (180 lines, Table S1, Figure 1B). Specifically, we recorded spontaneous mEPSPs and AP-evoked EPSCs after applying sub-saturating concentrations of the glutamate receptor (GluR) antagonist PhTX-433 (PhTX) for 10 minutes (20 μM; extracellular Ca^2+^ concentration, 1.5 mM). At WT NMJs, PhTX treatment significantly reduced mEPSP amplitude compared to untreated controls (Figure 1C, black and gray arrow), indicating GluR perturbation. By contrast, AP-evoked EPSC amplitudes were similar between PhTX-treated and untreated WT NMJs (Figure 1D, black and gray arrow). Together with a reduction in mEPSP amplitude, a similar EPSC amplitude suggests a homeostatic increase in neurotransmitter release after PhTX treatment, consistent with PHP ^5^. PhTX also reduced mean mEPSP amplitudes in the 180 transgenic or mutant lines (either presynaptic/neural RNAi expression, *elav*^*c155*^-*Gal4 > UAS-RNAi*; or mutations within the respective coding sequence, see Methods) compared to untreated WT controls (Figure 1C). Moreover, the mean EPSC amplitude of the majority of the tested lines did not differ significantly from the mean WT EPSC recorded at PhTX-treated NMJs (Figure 1D, white bars). The combination of a decrease in mEPSP amplitude and largely unchanged EPSC amplitude indicates that the majority of the tested lines likely display PHP. We also identified 21 transgenic or mutant lines with significantly smaller EPSC amplitudes compared to PhTX-treated WT NMJs, and two lines with increased EPSC amplitudes (Figure 1D, E, red data). These represent candidate mutations that may disrupt PHP. One of the mutant lines with significantly smaller EPSC amplitudes in the presence of PhTX was a previously described deletion of the gene *thin* (*CG15105*, *thin*^*ΔA*; 25^) (Figure 1E, filled red data). This gene was selected for further analysis.

### Presynaptic *thin* promotes rapid PHP expression

In the genetic screen, we compared synaptic transmission between a given genotype and WT controls in the presence of PhTX (Figure 1C-E). Hence, the small EPSC amplitude of *thin*^*ΔA*^ mutants seen after PhTX application could be either due to impaired PHP, or a defect in baseline synaptic transmission. To distinguish between these possibilities, we next quantified synaptic transmission in the absence and presence of PhTX in *thin*^*ΔA*^ mutants (Figure 2). Similar to WT controls, PhTX application significantly reduced mEPSC amplitude by ~40% in *thin*^*ΔA*^ mutants (Figure 2A, B), suggesting similar receptor impairment. At WT synapses, EPSC amplitudes were similar in the absence and presence of PhTX (Figure 2A and C). In combination with the decrease in mEPSC amplitude (Figure 2B), PhTX incubation increased quantal content (EPSC amplitude/mEPSC amplitude) in WT (Figure 2D), indicating homeostatic release potentiation. By contrast, PhTX treatment significantly reduced EPSC amplitudes in *thin*^*ΔA*^ mutants (Figure 2A, C) and did not increase quantal content (Figure 2D). These data indicate that *thin*^*ΔA*^ is required for acute PHP expression.

**Figure 2.**
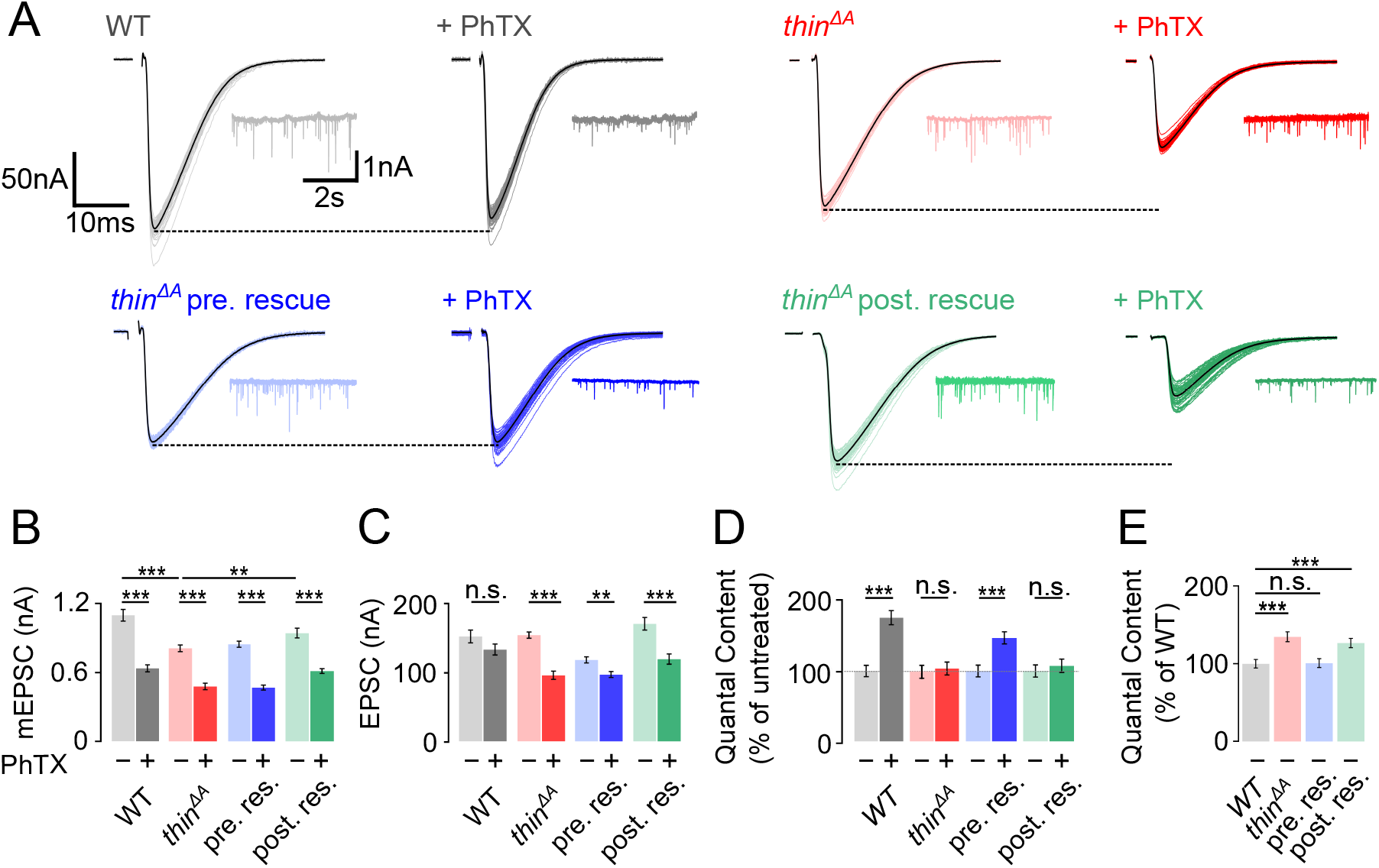
Homeostatic plasticity requires presynaptic *thin*. **A)** Representative EPSCs (individual sweeps and averages are shown in light colors and black, respectively), and mEPSCs (insets) of WT (gray), *thin*^*ΔA*^ mutants (red), presynaptic *thin* expression (*elav*^*C155*^-*Gal4* > *UAS-thin*, ‘*thin*^*ΔA*^ pre rescue’, blue), and postsynaptic *thin* expression (*24B-Gal4* > *UAS-thin*, ‘*thin*^*ΔA*^ post rescue’, green) in the *thin*^*ΔA*^ mutant background in the absence and presence of PhTX (‘+ PhTX’, darker colors). Stimulation artifacts were blanked for clarity. Note the decreased EPSC amplitudes at PhTX-treated *thin*^*ΔA*^ mutant NMJs and *thin*^*ΔA*^ post rescue NMJs, indicating impaired PHP. **B – E)** Mean mEPSC amplitudes (B), EPSC amplitudes (C), quantal content after PhTX treatment normalized to the respective untreated control, in the absence (‘−’) and presence (‘+’) of PhTX (D), and baseline quantal content of the indicated genotypes in the absence (‘−’) of PhTX normalized to WT (E). Note that PhTX did not enhance quantal content in *thin*^*ΔA*^ mutants, indicating impaired PHP. Also note the increased quantal content under baseline conditions in *thin*^*ΔA*^ mutants, suggesting increased release. Both phenotypes are restored upon presynaptic *thin* expression in the mutant background. Mean ± s.e.m.; n≥23 NMJs; *p < 0.05; **p < 0.001; ***p < 0.0001; n.s.: not significant; two-way ANOVA followed by Tukey’s post hoc test (B, C, D) and one-way ANOVA with Tukey’s multiple comparisons (E).

To test if presynaptic or postsynaptic *thin*^*ΔA*^ promotes PHP, we assessed PHP after presynaptic or postsynaptic expression of a *thin* transgene in the *thin*^*ΔA*^ mutant background. PhTX treatment significantly reduced mEPSC amplitudes after neural/presynaptic (*elav*^*c155*^-*Gal4*) or postsynaptic (*24B-Gal4*) expression of *thin* (*UAS-thin*) in *thin*^*ΔA*^ mutants (Figure 2A, B). While EPSC amplitudes were similar in the absence and presence of PhTX after presynaptic *thin* expression (‘presynaptic rescue’ or ‘pre. rescue’; Figure 2A, C, blue data), PhTX application significantly reduced EPSC amplitudes after postynaptic *thin* expression in the *thin*^*ΔA*^ mutant background (‘postsynaptic rescue’ or ‘post. rescue’; Figure 2A, C, green data). Together with the decrease in mEPSC amplitude, presynaptic, but not postsynaptic *thin* expression significantly enhanced quantal content in the *thin*^*ΔA*^ mutant background (Figure 2D). Thus, presynaptic, but not postsynaptic *thin* expression restores PHP in the *thin*^*ΔA*^ mutant background, implying a presynaptic role for *thin* in PHP.

We also noted a decrease in mEPSC amplitude in *thin*^*ΔA*^ mutants compared to WT in the absence of PhTX (Figure 2A, B), which is most likely due to impaired muscle architecture in *thin*^*ΔA*^ mutants ^25,26^. Postsynaptic, but not presynaptic *thin* expression, significantly increased mEPSC amplitudes towards WT levels in the *thin*^*ΔA*^ mutant background (Figure 2A, B), suggesting that postsynaptic *thin* is required for normal mEPSC amplitude levels. Furthermore, *thin*^*ΔA*^ mutants displayed a significant increase in quantal content compared to WT under baseline conditions in the absence of PhTX (Figure 2E), which was rescued by presynaptic, but not postsynaptic *thin* expression (Figure 2E). These data are consistent with the idea that presynaptic *thin* represses release under baseline conditions (see Figure 4, 7). By extension, the increased release under baseline conditions in *thin*^*ΔA*^ mutants may occlude PHP in response to receptor perturbation (see Discussion).

### *thin* promotes sustained PHP expression

PHP is not only expressed after acute pharmacological receptor perturbation, but also upon sustained genetic receptor impairment. At the *Drosophila* NMJ, genetic ablation of the GluRIIA subunit in *GluRIIA*^*SP16*^ mutants reduces quantal size and induces sustained PHP ^4^. To test if *thin* is required for sustained PHP expression, we generated recombinant flies carrying the *GluRIIA*^*SP16*^ and the *thin*^*ΔA*^ mutation (‘*GluRIIA*^*SP16*^, *thin*^*ΔA*^’). *GluRIIA*^*SP16*^ mutant NMJs displayed a strong decrease in mEPSC amplitude compared to WT (by ~50%; Figure 3A, B), which was accompanied by a significant increase in quantal content (Figure 3D) that restored EPSC amplitudes towards WT levels (Figure 3A, C), in line with previous work ^4^. By contrast, while mEPSC amplitudes were decreased by ~40% in *GluRIIA*^*SP16*^, *thin*^*ΔA*^ double mutants with regard to *thin*^*ΔA*^ mutants (Figure 3A, B), there was no increase in quantal content in *GluRIIA*^*SP16*^, *thin*^*ΔA*^ double mutants (Figure 3D), resulting in significantly smaller EPSC amplitudes than in *thin*^*ΔA*^ *mutants* (Figure 3A, C). Hence, *thin* is also necessary for sustained PHP expression, providing independent evidence for its role in homeostatic release regulation.

**Figure 3.**
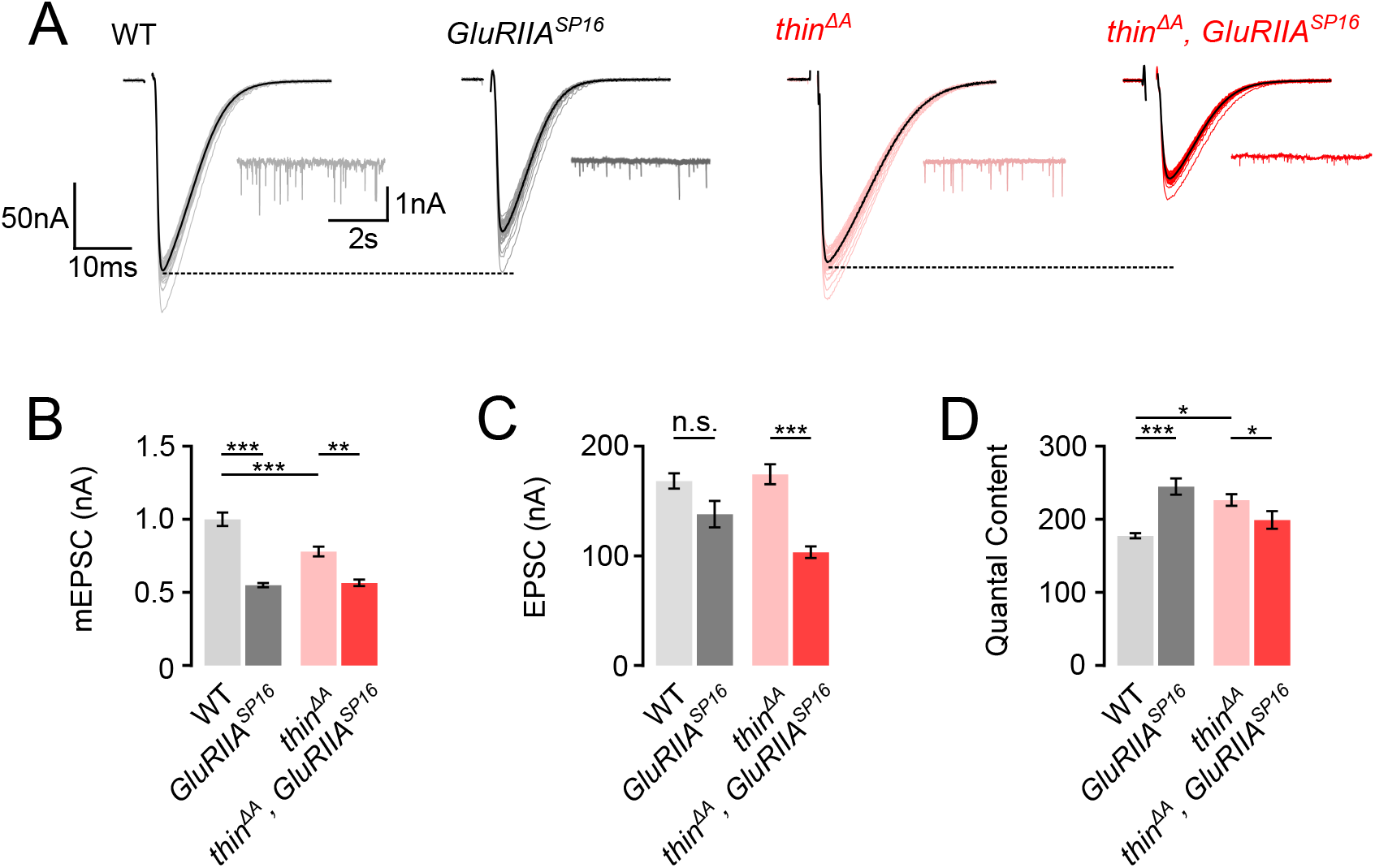
Sustained homeostasis is impaired in *thin* mutants. **A)** Representative EPSCs (individual sweeps and averages are shown in light colors and black, respectively), and mEPSCs (insets) of WT (gray), *GluRIIA*^*SP16*^ mutants (dark gray), *thin*^*ΔA*^ mutants (red), and *thin*^*ΔA*^, *GluRIIA*^*SP16*^ double mutants (dark red). Stimulation artifacts were blanked for clarity. **B – D)** Mean mEPSC amplitudes (B), EPSC amplitude (C), and quantal content (D) of the indicated genotypes. Note that there is no quantal content increase in *thin*^*ΔA*^, *GluRIIA*^*SP16*^ compared to *thin*^*ΔA*^, indicating impaired PHP. Mean ± s.e.m.; n≥13 NMJs; *p < 0.05; **p < 0.001; ***p < 0.0001; n.s.: not significant; Two-way ANOVA followed by Tukey’s post hoc test.

### *thin* negatively regulates release-ready vesicle number

Having established that *thin* is essential for acute and sustained PHP, we next explored the role of *thin* in the regulation of neurotransmitter release under baseline conditions. *thin*^*ΔA*^ mutants display increased neurotransmitter release in the absence of PhTX, and this increase in release is rescued by presynaptic *thin* expression (Figure 2).

To elucidate the mechanisms through which *thin* negatively modulatesrelease, we probed the size of the readily-releasable pool of synaptic vesicles (RRP) and neurotransmitter release probability (*p*_*r*_) after presynaptic *thin* perturbation (Figure 4). As the decreased mEPSC amplitude in *thin*^*ΔA*^ mutants may confound conclusions regarding presynaptic *thin* function (Figure 2), we focused our further analyses on the effects of presynaptic *thin*^*RNAi*^ expression (Figure 4). Presynaptic *thin*^*RNAi*^ expression (*elav*^*c155*^-*Gal4* > *UAS-thin*^*RNAi*^) significantly increased EPSC amplitudes (Figure 4A, C), with no significant effects on mEPSC amplitudes compared to controls (*elav*^*c155*^-*Gal4*/+; Figure 4A, B), suggesting that presynaptic *thin* represses release, consistent with the data obtained from *thin*^*ΔA*^ mutants (Figure 2).

**Figure 4.**
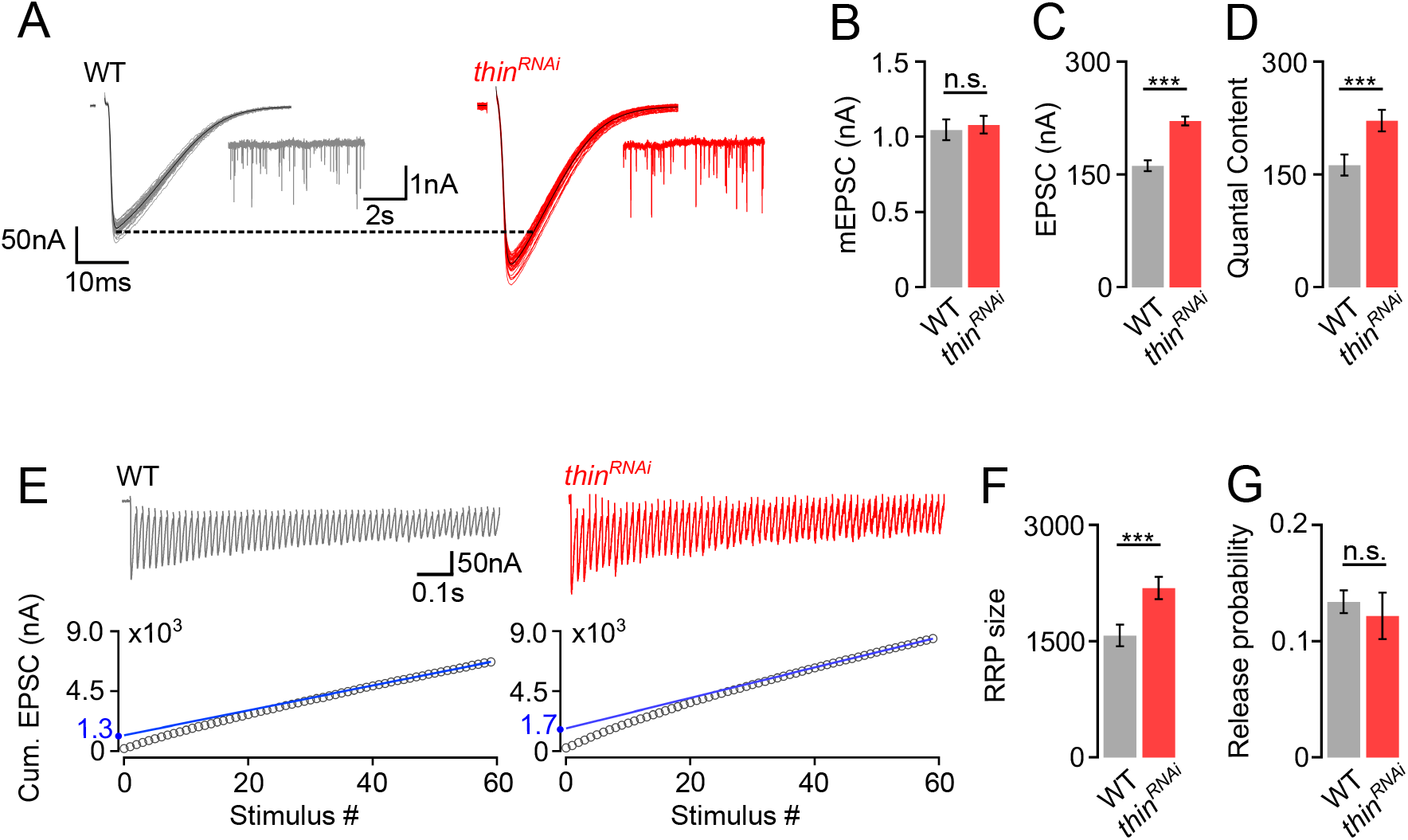
*thin* negatively regulates release-ready vesicle number. **A)** Representative EPSCs (individual sweeps and averages are shown in light colors and black, respectively), and mEPSCs (insets) of WT (gray) and presynaptic *thin*^*RNAi*^ (*elav*^*C155*^-*Gal4* > *UAS-thin*^*RNAi*^, red). **B – D)** Mean mEPSC amplitudes (B), EPSC amplitudes (C), and quantal content (D) of the indicated genotypes. **E)** Representative EPSC train (60 Hz, 60 stimuli, top) and cumulative EPSC amplitudes (‘cum. EPSC’, bottom) of WT and presynaptic *thin*^*RNAi*^. **F, G)** Mean readily-releasable vesicle pool (RRP) size (cum. EPSC/mEPSC) (F), and release probability (EPSC/cum. EPSC) (G) of the indicated genotypes. Note the increase in EPSC amplitude and RRP size in presynaptic *thin*^*RNAi*^. Mean ± s.e.m.; n≥22 NMJs; *p < 0.05; **p < 0.001; ***p < 0.0001; ns: not significant; Student’s *t*-test.

Next, we estimated RRP size using cumulative EPSC amplitude analysis during high-frequency stimulation (60 Hz; ^29,30^) (Figure 4E). This analysis revealed a significantly larger RRP size upon presynaptic *thin*^*RNAi*^ expression compared to controls (Figure 4E, F), implying that presynaptic *thin* negatively regulates RRP size. We then estimated *p*_*r*_ based on the ratio between the first EPSC amplitude of the stimulus train and the cumulative EPSC amplitude and observed no significant *p*_*r*_ differences between *thin*^*RNAi*^ and controls (Figure 4G). Thus, *thin* represses release by limiting the number of release-ready synaptic vesicles with largely unchanged *p*_*r*_.

### Altered NMJ development unlikely causes PHP defect in *thin* mutants

The PHP defect and the release enhancement under baseline conditions after presynaptic *thin* perturbation may arise from impaired synaptic development. To test this possibility, we investigated NMJ morphology in *thin* mutants (Figure 5). We confined the morphological analysis to NMJs lacking *thin* presynaptically (*thin^ΔA^*; *24BGal4>UAS-thin*; henceforth called ‘presynaptic *thin*^*ΔA*^ mutant’; Figure 5B), because ubiquitous loss of *thin* impairs muscle development ^25,26^. Immunostainings with an antibody detecting neuronal membrane (anti-horseradish peroxidase, ‘HRP’; ^31^) revealed a slight, but significant increase in HRP area in presynaptic *thin*^*ΔA*^ mutants compared to WT (Figure 5C), indicating a slight increase in NMJ size. Analysis of the active zone marker Bruchpilot (anti-Bruchpilot, ‘Brp’; ^32^) uncovered a significant increase in Brp puncta number per NMJ (Figure 5D), a slight increase in Brp density (Figure 5E), as well as a decrease in Brp intensity (Figure 5F) in presynaptic *thin*^*ΔA*^ mutants. In principle, these morphological changes could be related to the PHP defect or release enhancement seen in *thin*^*ΔA*^ mutants. However, postsynaptic *thin* expression in WT induced similar morphological alterations compared to *thin* overexpression in *thin*^*ΔA*^ mutants (presynaptic *thin*^*ΔA*^ mutants) (Figure S2), but neither impaired PHP nor enhanced release (Figure S2). These data suggest that the morphological changes seen in presynaptic *thin*^*ΔA*^ mutants are caused by postsynaptic *thin* expression rather than the *thin*^*ΔA*^ mutation, and that these morphological alterations do not block PHP or enhance release. Together, we conclude that the PHP defect and the increase in baseline synaptic transmission in *thin* mutants are unlikely caused by major changes in NMJ development.

**Figure 5.**
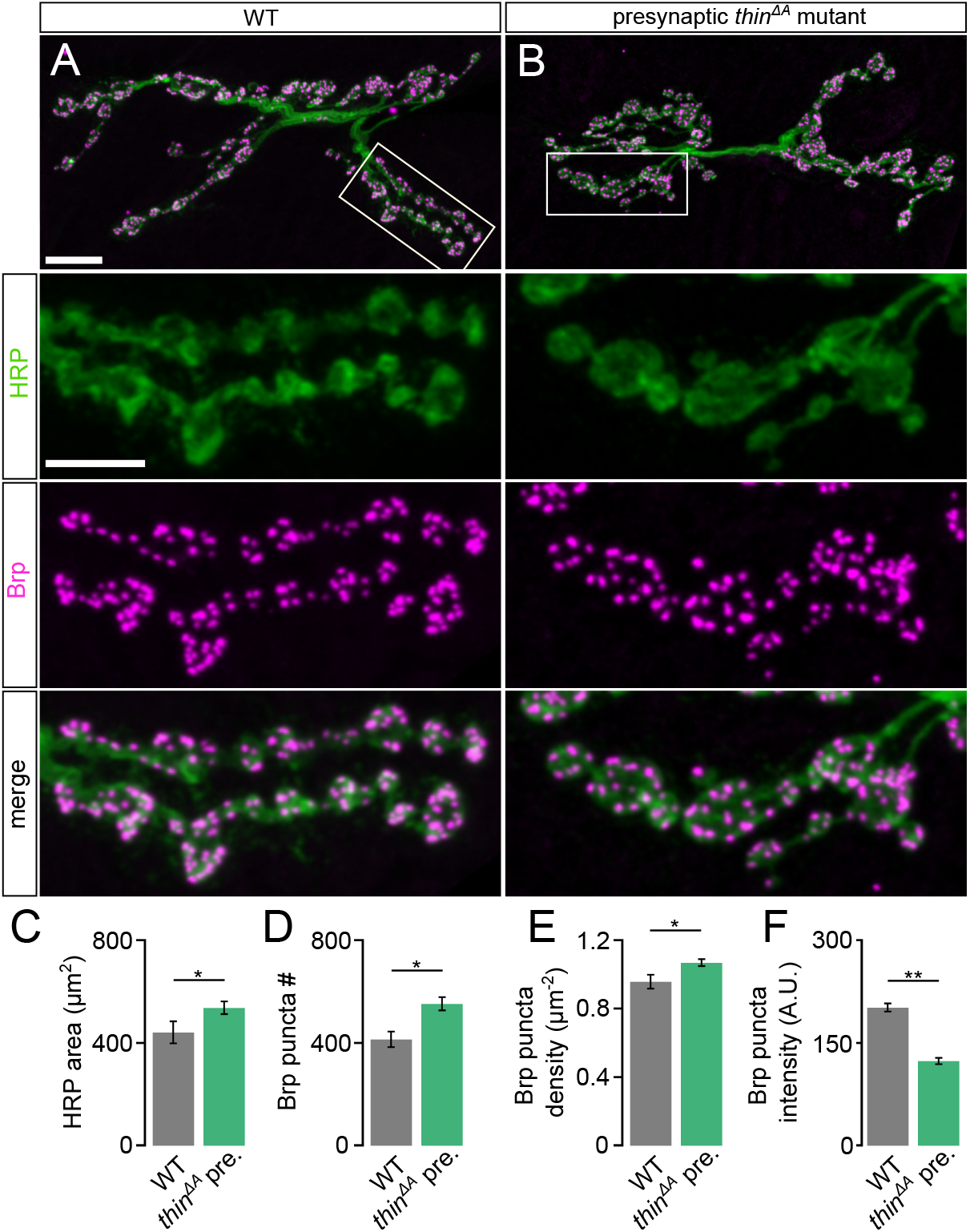
Altered NMJ development unlikely causes PHP defect in *thin* mutants. **A)** Maximum intensity projection of a WT NMJ, and **B)** an NMJ lacking *thin* presynaptically (*thin*^*ΔA*^; *24B-Gal4>UAS-thin* or ‘*thin*^*ΔA*^ pre.’ stained against the *Drosophila* neuronal membrane marker anti-HRP (‘HRP’) and the active-zone marker Bruchpilot (‘Brp’); scale bar, overview, 10 μm; inset, 5 μm. **C – F)** Mean HRP area (‘HRP area’, C), Brp puncta number per NMJ (‘Brp puncta #’, D), Brp puncta number/HRP area per NMJ (‘Brp puncta density’, E), Brp puncta fluorescence intensity (‘Brp puncta intensity’, F). Loss of presynaptic *thin* induces a slight, but significant increase in the total number of brp puncta. Mean ± s.e.m.; n≥10 NMJs; *p < 0.05; **p < 0.001; ***p < 0.0001; ns: not significant; Student’s t-test.

### Thin localizes in close proximity to Dysbindin and promotes Dysbindin degradation

Trim32, Thin’s human ortholog, ubiquitinates Dysbindin and targets it for degradation ^33^. *dysbindin*, in turn, is required for PHP at the *Drosophila* NMJ ^22^, and genetic evidence suggests that the UPS controls Dysbindin under baseline conditions and during PHP ^24^. We therefore explored the relationship between Thin and Dysbindin. First, we investigated the localization of Thin in relation to Dysbindin within synaptic boutons (Figure 6A). Previous studies suggest very low endogenous Dysbindin levels that preclude direct immunohistochemical analysis at the *Drosophila* NMJ ^22,24^. However, presynaptic expression of a fluorescently-tagged *dysbindin* transgene revealed that Dysbindin localizes in close proximity to synaptic vesicle markers ^22^ (Figure S3). The localization of fluorescently-tagged Dysbindin likely overlaps with the one of endogenous Dysbindin, as its presynaptic expression rescues the PHP defect in *dysbindin* mutants ^22^. The strong anti-Thin staining of *Drosophila* muscles makes it difficult to distinguish between presynaptic and postsynaptic Thin ^25^. This prompted us to analyse the localization of fluorescently-tagged Thin, which we expressed presynaptically (*elav*^*C155*^-*Gal4 > UAS-thin*^*mcherry*^). Presynaptic Thin^mcherry^ partially overlapped with fluorescently-tagged Dysbindin at confocal resolution (*elav*^*C155*^-*Gal4 > UAS-dysb*^*venus*^; Figure 6A, B). The localization of fluorescently-tagged Thin also likely overlaps with endogenous Thin, because presynaptic *thin* expression restores PHP and synaptic transmission in *thin* mutants (Figure 2). As indicated by the line profile across a bouton (Figure 6B), Dysbindin and Thin fluorescence intensity increased toward the bouton periphery (Figure 6B), similar to synaptic vesicle markers, such as synapsin (Figure S3B). At STED resolution, fluorescently-tagged Thin and Dysbindin appeared as distinct spots that partially overlapped (Figure 6C). Based on the close proximity between Dysbindin and synaptic vesicle markers ^22^, these data indicate that a fraction of Thin localizes in the close vicinity of Dysbindin and synaptic vesicles.

**Figure 6.**
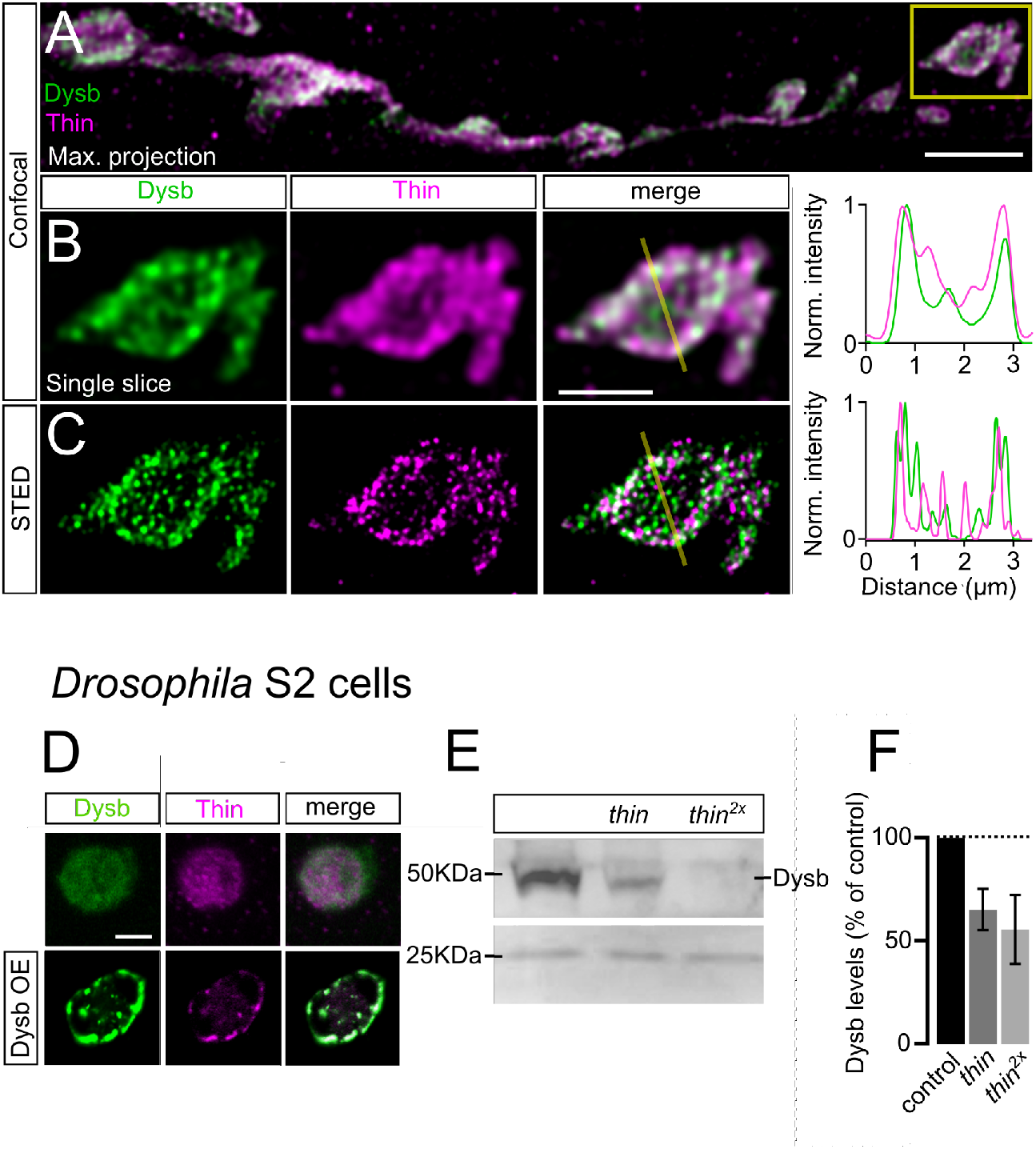
Thin localizes in close proximity to Dysbindin and promotes Dysbindin degradation. **A)** Confocal maximum intensity projection of a representative NMJ branch (muscle 6-7) after presynaptic co-expression (*elav*^*c155*^-*Gal4*) of venus-tagged Dysbindin (*UAS-dysb*^*venus*^, ‘Dysb’, green) and mCherry-tagged Thin (*UAS-thin*^*mcherry*^, ‘Thin’, magenta). **B)** Single plane of the synaptic bouton highlighted by the yellow square in (A) with corresponding line profile (*right*). The yellow line demarks the location of the line profile. **C)** gSTED image of the synaptic bouton shown in (B) with corresponding line profile (*right*). Note the partial overlap between Thin and Dysbindin at confocal and STED resolution. **D)** Confocal images (single planes) of *Drosophila* S2 cells stained with anti-Dysbindin (green) and anti-Thin (magenta) under control conditions (*top row*) and after *dysbindin* overexpression (*UAS-dysb*^*venus*^, *bottom row*). Note the concomitant redistribution of Dysbindin and Thin upon *dysbindin* overexpression. **E)** Representative Western blot of S2 cells transfected with *UAS-dysb*^*venus*^ and different levels of *UAS-thin*. **F)** Quantification of (E, n=5). Note the decrease in Thin levels upon *dysbindin* overexpression. Scale bar (A: 5μm), (B, C: 2μm), (D: 5μm).

Previous work in cultured human cells showed that Dysbindin ubiquitination by Thin’s human ortholog Trim32 induces Dysbindin degradation^33^. To test if this function is conserved in *Drosophila*, we used cultured *Drosophila* S2 cells. First, we probed the relationship between Thin and Dysbindin localization. Interestingly, while anti-Thin fluorescence was homogenously distributed within S2 cells under control conditions (Figure 6D, top), *dysbindin* (*dysb*^*venus*^) overexpression led to a redistribution of anti-Thin fluorescence into clusters that localized in close proximity to Dysbindin clusters (Figure 6D, bottom), suggesting a possible interaction between Thin and Dysbindin.

Next, we assessed whether Thin expression affects Dysbindin abundance in S2 cells by Western blot analysis. We observed a decrease in Dysb^venus^ levels upon co-expression of increasing Thin^mcherry^ levels (Figure 6E, F). Together, these data are consistent with the idea that Thin acts as an E3 ligase for Dysbindin in *Drosophila*, similar to Trim32 in humans ^33^.

### *thin* represses release through *dysbindin*

We next explored a possible genetic interaction between *thin* and *dysbindin* in the context of synaptic physiology. As *thin* and *dysbindin* mutants alone disrupt PHP, the analysis of double mutants would not be informative. We therefore investigated baseline synaptic transmission after presynaptic *thin*^*RNAi*^ expression in the *dysbindin* mutant background (Figure 7). Neither presynaptic *thin*^*RNAi*^ expression (*elav*^*c155*^-*Gal4* > *UAS-thin*^*RNAi*^) in the WT background, nor in the *dysb*^*1*^ mutant background affected mEPSC amplitude (Figure 7A, B). While presynaptic *thin*^*RNAi*^ expression enhanced EPSC amplitude and quantal content in the WT background (Figure 7C, D; see also Figure 4), presynaptic *thin*^*RNAi*^ expression did not change EPSC amplitude (Figure 7C) or quantal content (Figure 7D) in the *dysb*^*1*^ mutant background. These data provide genetic evidence that *thin* negatively controls release through *dysbindin* (Figure 7E).

**Figure 7.**
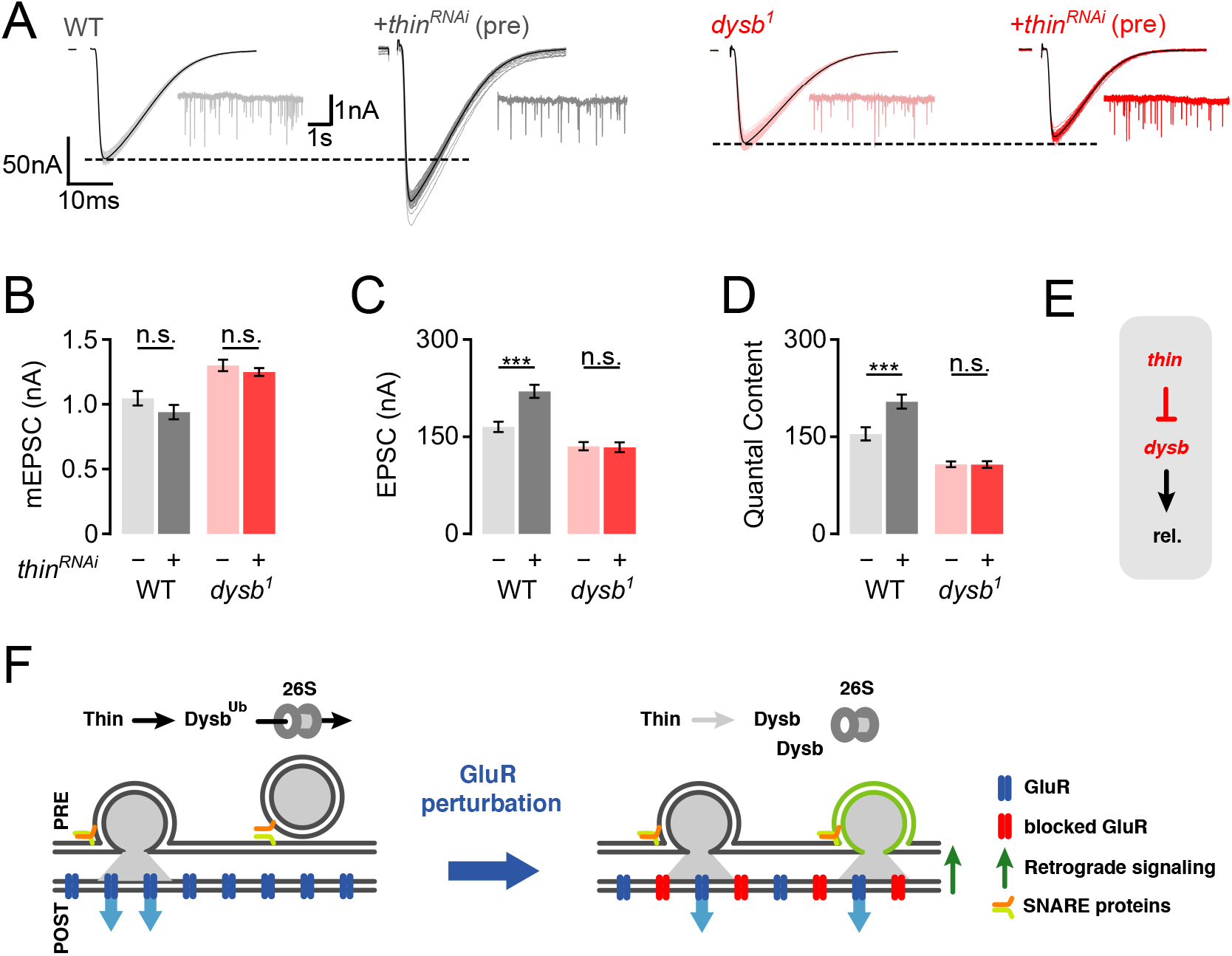
*thin* represses release through *dysbindin.* **A)** Representative EPSCs (individual sweeps and averages are shown in light colors and black, respectively), and mEPSCs (insets) of WT (gray) and presynaptic *thin*^*RNAi*^ (*elav*^*C155*^-*Gal4* > *UAS-thin*^*RNAi*^, dark gray), *dysb*^*1*^ mutants (light red), and presynaptic *thin*^*RNAi*^ in the *dysb*^*1*^ mutant background (*elav*^*C155*^-*Gal4* > *UAS-thin*^*RNAi*^, dark red). **B – D)** Mean mEPSC amplitudes (B), EPSC amplitudes (C), and quantal content (D) of the indicated genotypes. Note that presynaptic *thin*^*RNAi*^ expression increases EPSC amplitude and quantal content in WT, but not in *dysb*^*1*^ mutants. Mean ± s.e.m.; n≥13 cells; *p < 0.05; **p < 0.001; ***p < 0.0001; ns: not significant; two-way ANOVA followed by Tukey’s post hoc test. **E)** Working model: Our genetic data support a model in which *thin* controls neurotransmitter release (‘rel.’) through negative regulation of *dysbindin*. **F)** Emerging model: *Left:* Under baseline conditions, Thin ubiquitinates (‘Ub’) Dysbindin (‘Dysb’) and targets it for degradation by the 26S proteasome (‘26S’). *Right:* GluR perturbation (*red GluRs*) induces retrograde PHP signaling (green arrow), which decreases Thin-dependent Dysb degradation through an unknown pathway, thereby increasing Dysb levels and presynaptic release (green synaptic vesicle). Based on previous data, Dysbindin likely increases release by interacting with the SNARE complex through snapin and SNAP-25 (see Discussion).

## DISCUSSION

Employing an electrophysiology-based genetic screen targeting 157 E3 ligase-encoding genes at the *Drosophila* NMJ, we discovered that a mutation in the E3 ligase-encoding gene *thin* disrupts acute and sustained PHP. Presynaptic loss of *thin* led to increased release and RRP size, largely independent of gross synaptic morphological changes. Thin and Dysbindin localize in close proximity within synaptic boutons, and biochemical evidence suggests that Thin degrades Dysbindin *in vitro*. Finally, presynaptic *thin* perturbation did not enhance release in the *dysbindin* mutant background, providing genetic evidence that *thin* represses release through *dysbindin.*

As *thin* and *dysbindin* are required for PHP, these data are consistent with a model in which *thin* controls neurotransmitter release during PHP and under baseline conditions through *dysbindin* (Figure 7E, F).

Our study represents the first systematic investigation of E3 ligase function in the context of synaptic transmission. A considerable fraction of lines tested (11%) displayed a decrease in EPSC amplitude after PhTX treatment (Figure 1C-E). These E3 ligase-encoding genes may either be required for PHP and/or baseline synaptic transmission. Previous PHP screens in the same system identified PHP mutants with a success rate of ~3% ^22,23^. Thus, our data indicate that E3 ligase function either plays a special role in PHP and/or baseline synaptic transmission. As more transgenic or mutant lines exhibited a decrease in synaptic transmission, we conclude that the net effect of E3 ligases is to promote synaptic transmission at the *Drosophila* NMJ. Given the evolutionary conservation of most E3 ligases tested in this study (Figure 1, Table S1), the results of our screen likely allow predicting the role of the tested E3 ligases in neurotransmitter release regulation in other systems.

Previous studies linked E3 ligases to synaptic development and synaptic function at the *Drosophila* NMJ ^34,35^. For instance, the E3 ligase *highwire* (*hiw*) restrains synaptic growth and promotes evoked synaptic transmission at the *Drosophila* NMJ ^34^. Similarly, the deubiquitinating protease *fat facets* represses synaptic growth and enhances synaptic transmission ^36^. Although different molecular pathways have been implicated in *hiw*-dependent regulation of synaptic growth and function ^16^, it is generally difficult to disentangle effects on synaptic morphology from synaptic function. *thin* and its mammalian ortholog *Trim32* are required for maintaining the cytoarchitecture of muscle cells ^25,37,38^. Hence, the changes in synaptic transmission described in the present study may be a secondary consequence of impaired muscle structure. However, presynaptic *thin* expression in the *thin* mutant background restored synaptic function under baseline conditions and during homeostatic plasticity (Figure 2). Conversely, while postsynaptic *thin* expression largely rescued the defects in muscle morphology in *thin* mutants, the defects in synaptic function persisted. These genetic data suggest that the impairment of baseline synaptic transmission and homeostatic plasticity in *thin* mutants is unlikely caused by muscular dystrophy. We also noted a slight increase in Brp number at presynaptic *thin* mutant NMJs (Figure 5), indicating increased active-zone number. In principle, this increase in active-zone number may underlie the increase in neurotransmitter release after presynaptic loss of *thin*. However, postsynaptic *thin* overexpression increased Brp number in WT, but did neither affect baseline synaptic transmission, nor PHP (Figure S2). Hence, our results suggest that presynaptic *thin* regulates neurotransmitter release under baseline conditions and during homeostatic plasticity largely independent of changes in synaptic morphology.

We revealed that presynaptic *thin* perturbation results in enhanced neurotransmitter release (Figure 2, 4, 7), indicating that the E3 ligase Thin represses neurotransmitter release under baseline conditions. Notably, there are just a few molecules that have been implicated in repressing neurotransmitter release, such as the SNARE-interacting protein tomosyn ^39,40^, or the RhoGAP crossveinless-c ^41^. How could the E3 ligase Thin oppose neurotransmitter release? We discovered that *dysbindin* is required for the increase in release induced by presynaptic *thin* perturbation (Figure 7). Moreover, we revealed that Thin localizes in close proximity to Dysbindin in synaptic boutons (Figure 6), and that Thin degrades Dysbindin *in vitro* (Figure 6), similar to its mammalian ortholog Trim32 ^33^. At the *Drosophila* NMJ, 26S-proteasomes are transported to presynaptic boutons ^42^, where they degrade proteins on the minute time scale ^24,43^. Previous genetic data suggest a positive correlation between Dysbindin levels and neurotransmitter release ^22,24^, and there is genetic evidence for rapid, UPS-dependent degradation of Dysbindin at the *Drosophila* NMJ ^24^. In combination with these previous observations, our data are consistent with the idea that Thin opposes release by acting on Dysbindin. Although the low abundance of endogenous Dysbindin at the *Drosophila* NMJ precludes direct analysis of Dysbindin levels ^22^, we speculate that Thin decreases Dysbindin abundance by targeting it for degradation. Alternatively, Thin may modulate Dysbindin function through mono-ubiquitination. Genetic data suggest that Dysbindin interacts with the SNARE protein SNAP25 through Snapin ^44^. Hence, Thin-dependent regulation of Dysbindin may modulate release via Dysbindin’s interaction with the SNARE complex.

Our study revealed a crucial role for *thin* in PHP. How does the increase in neurotransmitter release in *thin* mutants under baseline conditions relate to the PHP defect? The relative increase in release during PHP of WT synapses exceeds the increase in release in *thin* mutants under baseline conditions. Thus, although we cannot exclude that PHP is solely occluded by enhanced baseline release in *thin* mutants, we consider this scenario unlikely. PHP is blocked by acute pharmacological, or prolonged genetic proteasome perturbation at *Drosophila* NMJ ^24^. Moreover, PHP at this synapse requires *dysbindin* ^22^, and genetic data suggest UPS-dependent control of a Dysbindin-sensitive vesicle pool during PHP ^24^. Based on our finding that *thin* is required for acute and sustained PHP expression (Figures 2 and 3), and the links between *thin* und *dysbindin* in the context of release modulation outlined above, we propose a model in which Thin-dependent ubiquitination of Dysbindin is decreased during PHP (Figure 7F). Given the positive correlation between Dysbindin levels and release ^24,44^, the resulting increase in Dysbindin abundance would potentiate release. Further work is needed to test how Thin is regulated during PHP. Thin is the first E3 ubiquitin ligase linked to homeostatic regulation of neurotransmitter release. Interestingly, a recent study revealed a postsynaptic role for Insomniac, a putative adaptor of the Cullin-3 ubiquitin ligase complex, in PHP at the *Drosophila* NMJ ^45^, suggesting a key function of the UPS in both synaptic compartments during PHP at this synapse.

Trim32, the human ortholog of *thin*, is required for synaptic down-scaling in cultured hippocampal rat neurons ^46^, as well as long-term potentiation in hippocampal mouse slices ^47^, implying a broader role of this E3 ubiquitin ligase in synaptic plasticity. Trim32 has been implicated in various neurological disorders, such as depression ^48^, Alzheimer’s Disease ^49^, Autism Spectrum Disorder ^48,50^, or attention deficit hyperactivity disorder ^51^. It will be exciting to explore potential links between Trim32-dependent control of synaptic homeostasis and these disorders in the future.

## METHODS

### Fly stocks and genetics

*Drosophila* stocks were maintained at 21°C – 25°C on normal food. The *w*^*1118*^ strain was used as the wild-type (WT) control. *GluRIIA*^*SP16*^ mutants ^4^ and *dysbindin*^*1*^ mutants ^22^ were a kind gift from Graeme Davis’ lab. *thin*^*ΔA*^ mutants and *UAS-abba* transgenic flies, now referred to as *UAS-thin* ^25^, were a generous gift from Erika Geisbrecht. For pan-neuronal expression, the *elav*^*c155*^-*Gal4* (on the X chromosome) driver line was used and analysis was restricted to male larvae. For expression in muscle cells, the *24B-Gal4* driver line was used. Both driver lines were obtained from the Bloomington Drosophila Stock Center (Bloomington, IN, USA). Standard second and third chromosome balancer lines (Bloomington) and genetic strategies were used for all crosses and for maintaining the mutant lines. For the generation of transgenic flies carrying *UAS-thin::mcherry*, constructs based on the pUAST-attB vector backbone were injected into the ZP-attP-86Fb fly line harboring a landing site on the third chromosome according to standard procedures ^52^.

### Cell culture and transfection

Schneider S2 cells were cultivated in standard Schneider’s *Drosophila* medium (Gibco™) containing 10% fetal calf serum and 5% Penicilin/Streptomycin at 25 °C. For immunohistochemistry and microscopy cells, were plated on cover slips in 12 well plates with 80% density and transfected with 1.5 μg (total) vector DNA using FuGENE^®^ HD Transfection Reagent according to the standard protocol. Vectors used were: pMT-Gal4 (Addgene), pUAS-thin-mCherry and pUAS-venus-dysbindin (Dion Dickman). 24 h after plating, CuSO4 (0.5 mM) was added to the culture for 24 h to induce the expression of the pMT vector driving Gal4, which in turn drives transcription of UAS constructs.

### Plasmid construction

All plasmids were generated by standard restriction enzyme ligation. For the pUAS_attB_mCherry_thin vector, mCherry was cloned into pUAS_attB (Addgene) via EcoRI/NotI using the following primers (fw: 5’-ctcggcgcgccaATGGTGAGCAAGGGC GAGGAG-3’, rev: 5’-cgcggtaccttaCTTGTACAGCTCGTCCA TGCCGC-3’). *thin* was amplified from *Drosophila* cDNA by PCR using the following primers (fw : CGGAATTCATGGAGCAATTCGAGCAGCTGTTGACG, rev: CGTCTAGAATGAAGACTTGGACGCGGTGATTCTCTCG) and then cloned into the pUAS-attB-mcherry vector via NotI/XbaI. Correct cloning was confirmed by sequencing of all final vectors.

### Electrophysiology

Electrophysiological recordings were made from third-instar larvae at the wandering stage. Larvae were dissected and sharp-electrode recordings were made from muscle 6 in abdominal segments 3 and 4 using an Axoclamp 900A amplifier (Molecular Devices). The extracellular HL3 saline contained (in mM): 70 NaCl, 5 KCl, 10 MgCl_2_, 10 Na-Hepes, 115 sucrose, 5 trehalose, 5 HEPES, 1.5 CaCl_2_. To induce PHP, larvae were incubated with 20 μM PhTX-433 (Cat # sc-255421, Santa Cruz Biotechnology) for 10 min at room temperature after partial dissection (see ^5^). AP-evoked EPSCs were induced by stimulating hemi-segmental nerves with single APs (0.3 ms stimulus duration, 0.3 Hz), and recorded with a combination of a HS-9A x10 and a HS-9A x0.1 headstage (Molecular Devices) in two-electrode voltage clamp (TEVC) mode. mEPSPs and mEPSCs were recorded with one or two HS-9A x0.1 headstage(s) (Molecular Devices), respectively. Muscle cells were clamped to a membrane potential of −65 mV for EPSCs and −100 mV for mEPSCs to increase the signal-to-noise ratio. A total of 50 EPSCs were averaged to obtain the mean EPSC amplitude for each NMJ. RRP size was calculated by the method of cumulative EPSC amplitudes ^53^. NMJs were stimulated with 60-Hz trains (60 stimuli, 5 trains per cell), and the cumulative EPSC amplitude was obtained by back-extrapolating a linear fit to the last 15 cumulative EPSC amplitude values of the 60-Hz train to time zero.

### Immunohistochemistry and microscopy

#### Drosophila NMJ

Third-instar larval preparations were fixed for 3 min with Bouin’s fixative (100%, Sigma-Aldrich, HT-10132) for confocal microscopy, or 100% ice-cold Ethanol for 10 min for STED microscopy. Preparations were washed thoroughly with PBS containing 0.1% Triton X-100. After washing, preparations were blocked with 3% normal goat serum in PBS containing 0.1% Triton X-100. Incubation with the primary antibody was done at 4 °C on a rotating platform overnight.

The following antibodies and dilutions were used for NMJ stainings: (*Primary*) anti-Bruchpilot (nc82, mouse, DSHB, AB_2314866, 1:100), anti-GFP (rabbit, Thermo Fisher Scientific, G10362, 1:500), anti-GFP (mouse, Thermo Fisher Scientific, A-11120, 1:500), anti-DsRed (rabbit, Clontech, sc-390909, 1:500), anti-SYNORF1 (Synapsin, 3C11, mouse, DSHB, AB_528479, 1:250), anti-HRP Alexa-Fluor 647 (goat, Jackson ImmunoResearch 123–605–021, 1:200). For confocal microscopy Alexa-Fluor anti-mouse 488 (Thermo Fisher Scientific; 1:500) and Alexa Fluor anti-guinea pig 555 (Thermo Fisher Scientific; 1:400) were applied overnight at 4°C on a rotating platform. For gSTED microscopy (Figure 6,S3) the following secondary antibodies (1:100) were applied for 2 h at room temperature on a rotating platform: Atto 594 (anti-mouse, Sigma-Aldrich, 76085) and Abberior STAR 635 P (anti-rabbit, Abberior, 53399). Experimental groups of a given experiment were processed in parallel in the same tube. Preparations were mounted onto slides with ProLong Gold (Life Technologies, P36930).

#### S2 cell culture

S2 cells grown on coverslips were washed with PBST (PBS + 0.1% TritonX-100) and fixed with 10% PFA for 10 min. After washing three times with PBST, preparations were blocked with 5% normal goat serum in PBST for 30 min. Incubation with primary antibody was done at RT on a rotating platform for 2 h. The following antibodies were used for S2 cell stainings: anti-thin (guineapig, gift from Erika R. Geisbrecht, 1:200), anti-dysbindin (mouse, gift from Dion Dickman, 1:400). After washing three times with PBST, cells were incubated with secondary antibodies Alexa Fluor anti-guinea pig 555 and Alexa Fluor anti-mouse 488 (Thermo Fisher Scientific; 1:400) at RT on a rotating platform for 2 h. Cover slips were mounted onto slides with ProLong Gold (Life Technologies, P36930) after three PBST washes.

#### Confocal and gSTED microscopy

Images were acquired with an inverse Leica TCS SP8 STED 3X microscope (Leica Microsystems, Germany) of the University of Zurich Center for Microscopy and Image Analysis. Excitation light (580nm or 640nm) of a flexible white light laser was focused onto the specimen using a 100x objective (HC PL APO 1.40 NA Oil STED WHITE; Leica Microsystems, Germany) with immersion oil conforming to ISO 8036 with a diffraction index of n=1.5180 (Leica Microsystems, Germany). For gSTED imaging, the flexible white light laser was combined with a 775 nm STED depletion laser. Emitted light was detected with two HyD detectors in photon counting mode (Leica Microsystems, Germany). Pixel size was 20 × 20 nm and z-stacks were acquired with a step size of 120 nm. For STED imaging, we used time-gated single photon detection (empirical adjustment within a fluorescence lifetime interval from 0.7 to 6.0 ns). Pixel size was 20 × 20 nm and z-stacks were acquired with a step size of 120 or 130 nm. Line accumulation was set to 1 and 6 for confocal and STED imaging, respectively. Images were acquired with LAS X software (Leica Application Suite X, version 2.0; Leica Microsystems, Germany). Experimental groups were imaged side-by-side with identical settings.

Images were processed and deconvolved with Huygens Professional (Huygens compute engine 17.04, Scientific Volume Imaging B.V., Netherlands). In brief, the “automatic background detection” tool (radius=0.7μm), and the “auto stabilize” feature were used to correct for background and lateral drift. Images were deconvolved using the Good’s roughness Maximum Likelihood algorithm with default parameter settings (maximum iterations: 10; signal to noise ratio: 7 for STED and 15 for confocal; quality threshold: 0.003).

### Western blot

Transfected cells in 12-well-plates were washed with PBS and lysed by adding 50 μl of RIPA buffer (50 mM Tris, pH 8.0, 150 mM NaCl, 1% Nonidet P-40, 0.5% deoxycholate, 0.1% SDS, 0.2 mM sodium vanadate, 10 mM NaF, 0.4 mM EDTA, 10% glycerol) containing protease inhibitors (cOmplete™, Mini, EDTA-free Protease Inhibitor Cocktail, Sigma) for 30 min on ice. The lysates were sonified three times for 1 min and boiled for 5 min in SDS-sample buffer containing 5% ß-Mercaptoethanol. Samples were separated on acrylamide gels using SDS-PAGE, then transferred to nitrocellulose membranes (Amersham Hibond GE healthcare). After blocking in 5% milk in PBST, membranes were incubated in the following primary antibodies: anti-GFP (rabbit, Thermo Fisher Scientific, G10362; 1:500), anti-DsRed (rabbit, Clontech, sc-390909, 1:500) in blocking solution overnight. Horseradish peroxidase–conjugated secondary Abs (anti-mouse-HRP and anti-rabbit-HRP 1:2000 in blocking solution) were applied to membranes for 2 h. Detection was performed using ECL Reagent (GE Healthcare, Chicago, IL, USA).

### Data analysis

Electrophysiology data were acquired with Clampex (Axon CNS, Molecular Devices) and analysed with custom-written routines in Igor Pro (Wavemetrics). For the genetic screen data, mEPSPs were detected with a template matching algorithm implemented in Neuromatic (Rothman & Silver, 2018) running in Igor Pro (Wavemetrics). The average mEPSP amplitude was calculated from all detected events in a recording after visual inspection for false positives. For the rest of the data, mEPSC data were analysed using routines written with scientific python libraries, including numpy, scipy, IPython and neo ^54^, and mEPSCs were detected using an implementation of a template-matching algorithm ^55^.

Microscopy images were analysed using custom-written routines in ImageJ (version 1.51n, National Institutes of Health, USA). Brp quantification was performed as follows: First, individual Brp puncta were isolated by segmenting binary fluorescence intensity threshold masks (15% or 35% of the maximum intensity value) of background corrected (rolling ball, radius = 1 μm) and filtered (3 × 3 median) maximum intensity projection images. The number of Brp objects in the mask served as a proxy for AZ number, and was normalized to the area of the HRP mask (binary mask, 15% or 35% of the maximum intensity value). Average Brp-intensity values were calculated for each Brp punctum from background-corrected, unfiltered maximum intensity projection images.

Statistical analyses were done using RStudio Team (2021). RStudio: Integrated Development Environment for R. RStudio, PBC, Boston, MA. For more than two factors, we used two-way ANOVA followed by Tukey’s post hoc test to correct for multiple comparisons between genotypes and conditions. For one factor with more than two groups, one-way ANOVA with Tukey’s multiple comparisons was performed. Two-sided Student’s t-tests or nonparametric Mann-Whitney U tests were used for comparison between two groups after a Shapiro-Wilk test and a Levene’s test. Statistical significance was set to 0.05 (*), 0.01 (**) and 0.001 (***). Power analysis was performed using the pwr-package of Rstudio to estimate the minimum sample size for a power above ≥0.8 and a significance level of 0.05 for two-sided Student’s t tests or Mann-Whithiney U tests. Data are given as mean ± s.e.m.

Figures were assembled using GIMP (The GIMP team, 2.8.10, www.gimp.org) and Inkscape (Inkscape project, 0.92.2. http://www.inkscape.org).

## Acknowledgement

We are grateful to members of the Müller lab for helpful discussions and critical comments on the manuscript. We thank Dr. Damian Szklarczyk for help with STRING-based protein-protein interaction analysis used for prioritization of E3 ligases.

## Author Contributions

MB-C, KS and MM conceptualized and designed experiments. MB-C and KS conducted research and analysed data. MB-C, KS and MM interpreted data. MM and MB-C wrote the manuscript.

## Data Availability

All data needed to evaluate the conclusions in the paper are present in the paper and the Supplementary Materials and are available upon reasonable request.

