## Supplementary Material for "The E3 ligase Thin controls homeostatic plasticity through neurotransmitter release repression"

This PDF includes:

Supplementary Figures 1-3

Supplementary Table 1

### SUPPLEMENTARY INFORMATION

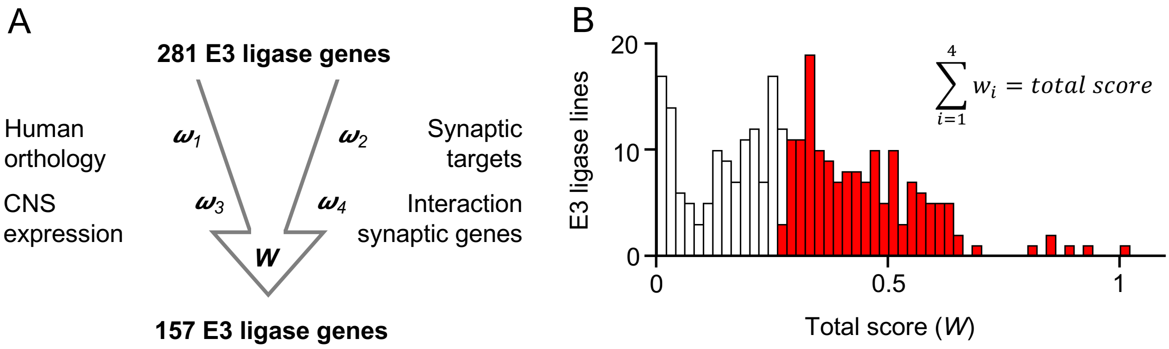


**Supplementary Figure 1. Generation and prioritization of the E3 ligase-encoding gene list for *Drosophila melanogaster***

**A)** Flow chart describing the prioritization process of the E3 list. *Generation:* First, we used the Gene Ontology (GO) search of Flybase [56] to identify genes annotated to encode for proteins with E3 ubiquitin ligase domains within the *Drosophila melanogaster* genome. This yielded an initial list of 221 genes, including confirmed and putative E3 ligase-encoding genes, similar to previous estimates [27]. Next, we added genes encoding domains contributing to the formation of the E3 complex, including the F-box domain, the Cullin domain, the N-recognin domain, the SKP1 domain, and the U-box domain. Subsequently, we searched human E3 ligase-encoding genes [57] for *Drosophila melanogaster* orthologs using the *Drosophila* RNAi Screening Center Integrative Ortholog Prediction Tool (DIOPT; version 8.0; http://www.flyrnai.org/diopt) [58]. In total, this approach identified 281 putative E3 ligase-encoding genes in the *Drosophila melanogaster* genome. *Prioritization:* We used a combination of four different criteria to create a score in order to prioritize the E3 list for screening the most relevant candidates (Normalized to max.). First, we prioritized for evolutionary conservation according to the overall DIOPT score of each putative E3 ligase-encoding gene with regard to its human ortholog (Hu et al., 2011) (score 1, "$\omega_1$"). Second, we prioritized for genes with predicted central nervous system expression based on transcriptomics data from modENCODE [59] and FlyAtlas [60] ($\omega_2$). Third, we prioritized for genes encoding for proteins predicted to interact with synaptic proteins. In short, we created a literature based list of known synaptic genes and calculated an interaction probability between each putative E3 ligase-encoding gene and all synaptic genes using STRING [61] ($\omega_3$). Fourth, we considered the probability of synaptic function predicted by machine-learning based analysis of transcriptomics data [62,63] ($\omega_4$). The list of putative E3 ligase-encoding genes was sorted according to the sum of the four scores for each gene. **B)** Distribution of the total score (summed weights) for all putative E3 ligase-encoding genes. The red bars indicate the lines selected for analysis (arbitrary threshold). In addition, we added to our screen genes encoding E3 ligases with known targets implicated in synaptic transmission and synaptic plasticity based on previously published data. Altogether, we tested 157 out of the 281 genes.

.

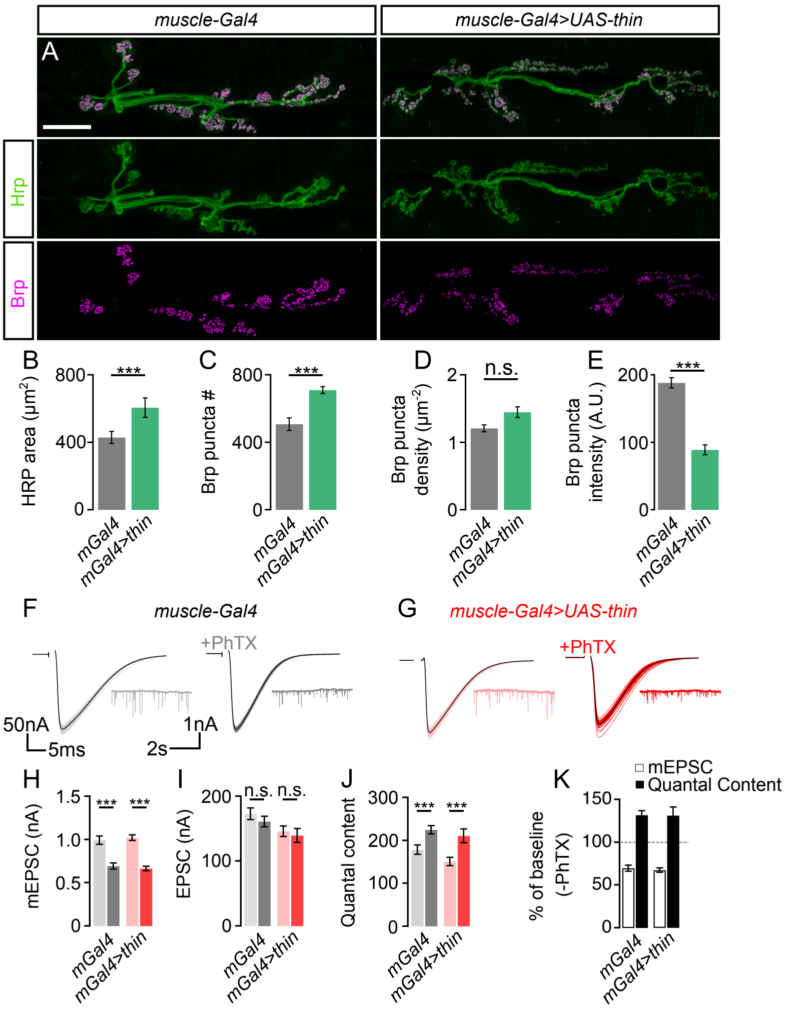


**Supplementary Figure 2. Postsynaptic *thin* overexpression increases active-zone number without affecting PHP**

**A)** Maximum intensity projection of a control NMJ (*24B-Gal4*/+, *'muscle Gal4'*, *left*) and an NMJ overexpressing *thin* in the muscle *24B-Gal4 > UAS-thin*, *right*) stained against the *Drosophila* neuronal membrane marker anti-HRP ('HRP') and the active-zone marker Bruchpilot ('Brp'; scale bar, 10 µm). **B – E)** Mean HRP area ('HRP area', B), Brp puncta number per NMJ ('Brp puncta #', C), Brp puncta number/HRP area per NMJ ('Brp puncta density', D), Brp puncta fluorescence intensity ('Brp puncta intensity', E). Postsynaptic *thin* overexpression induces an increase in neuronal area and Brp puncta number, as well as a decrease in Brp fluorescence intensity, similar to NMJs lacking *thin* presynaptically (*thin*$^{\Delta A}$; *24B-Gal4>UAS-thin*; see figure 5). Mean ± s.e.m.; (*24B-Gal4*/+: n≥10 NMJs; *24B-Gal4 > UAS-thin*: n≥10 NMJs). **F)** Representative EPSCs (individual sweeps and averages are shown in light colors and black, respectively), and mEPSCs (insets) of a control NMJ (*24B-Gal4*/+, *'muscle Gal4'*) in the absence (light gray) and presence (dark gray) of PhTX ('+PhTX', dark gray). Stimulation artifacts were blanked for clarity. **G)** Same as in F for an NMJ overexpressing *thin* in the muscle (*24B-Gal4 > UAS-thin*) **H – J)** Mean mEPSC amplitudes (H), EPSC amplitudes (**I**), and quantal content (J) of control (*24B-Gal4*/+, *'mGal4'*, gray) and postsynaptic *thin* overexpression (*24B-Gal4 > UAS-thin*, '*mGal4 > thin'*, red) in the absence (light colors) and presence of PhTX (dark colors). K) mEPSC amplitude (white) and quantal content (black) in the presence of PhTX normalized to control (without PhTX) of the indicated genotypes. Postsynaptic *thin* overexpression increases quantal content, indicating PHP expression. In combination with the morphology data (A – E), this implies that the PHP defect of NMJs lacking *thin* presynaptically (*thin*$^{\Delta A}$; *24B-Gal4>UAS-thin*; see figure 5) is unlikely due to an increase in Brp puncta number. Mean ± s.e.m.; (*24B-Gal4*/+: n=17 NMJs; *24B-Gal4 > UAS-thin*: 18 NMJs); *p < 0.05; **p < 0.001; ***p < 0.0001; n.s.: not significant; Student's t-test for pairwise comparison between control (*24B-Gal4*/+) and postsynaptic *thin* overexpression (*24B-Gal4>UAS-thin*).

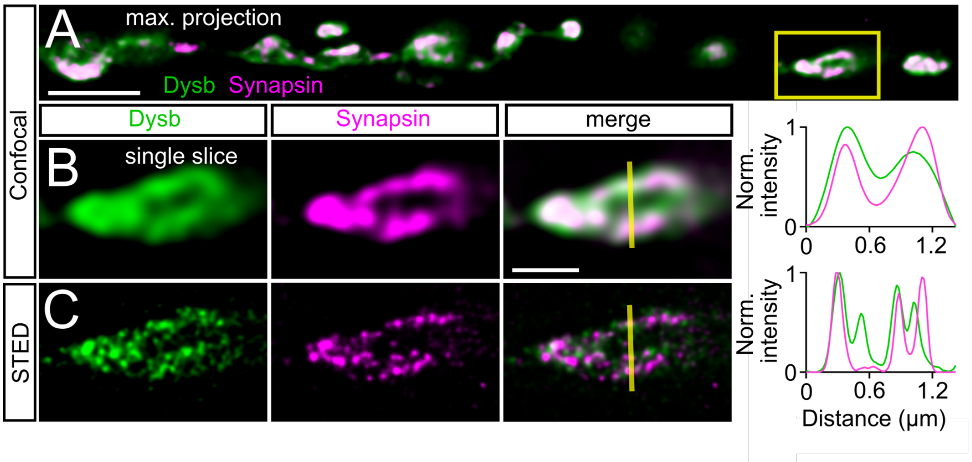


**Supplementary Figure 3. Dysbindin and Synapsin distribute in the periphery of synaptic boutons. A)** Confocal maximum intensity projection of a representative NMJ branch (muscle 6-7) after presynaptic expression (*elav*$^{c155}$-*Gal4*) of venus-tagged Dysbindin (*UAS-dysb*$^{venus}$) stained with anti-GFP (green, 'Dysb') and anti-Synapsin (magenta, 'Synapsin'). **B)** Single slice of the synaptic bouton highlighted by the yellow square in (A) with corresponding line profile (*right*). The yellow line demarks the location of the line profile. **C)** gSTED image of the synaptic bouton shown in (B) with corresponding line profile (*right*). Note the partial overlap between Dysbindin and the synaptic vesicle marker Synapsin at confocal and STED resolution. bar (A: 4µm), (B, C: 1µm).

**Supplementary Table S1**

| Gene name | EPSC (nA) | mEPSP (mV) |
|---|---|---|
| WT | -173.15 ± 4.8 | 0.65 ± 0.016 |
| CG12084 | -184.72 ± 22.8 | 0.43 ± 0.062 |
| Cand1 | -147.27 ± 9.1 | 0.54 ± 0.036 |
| rocl1a | -112.9 ± 6.3 | 0.54 ± 0.032 |
| phyl | -172.08 ± 21.5 | 0.41 ± 0.04 |
| Iap2 | -153.62 ± 10.1 | 0.51 ± 0.035 |
| Iap1 | -153.89 ± 17.5 | 0.89 ± 0.071 |
| CG3356 | -114.12 ± 19.6 | 0.34 ± 0.025 |
| tn | -99.21 ± 10.5 | 0.48 ± 0.012 |
| Ube3a line 1 | -151.47 ± 23.8 | 0.55 ± 0.078 |
| Ube3a line 2 | -96.66 ± 11.6 | 0.44 ± 0.031 |
| dbo | -195.01 ± 8.8 | 0.53 ± 0.015 |
| CG10080 | -131.63 ± 5.5 | 0.77 ± 0.132 |
| Cul1 line 1 | -138.96 ± 17.7 | 0.48 ± 0.019 |
| Cul1 line 2 | -100.95 ± 17.7 | 0.48 ± 0.034 |
| ctrip | -102.87 ± 12 | 0.46 ± 0.072 |
| Bre1 | -170.19 ± 30.4 | 0.7 ± 0.072 |
| Ago | -140.79 ± 11.3 | 0.63 ± 0.077 |
| msl-1 | -155.97 ± 13.5 | 0.95 ± 0.284 |
| Ebi | -130.77 ± 17.2 | 0.44 ± 0.055 |
| Nedd4 | -132.51 ± 4.8 | 0.5 ± 0.057 |
| Sina | -183.58 ± 16 | 0.53 ± 0.031 |
| FBXO11 | -173.24 ± 13.4 | 0.49 ± 0.012 |
| neur | -170.22 ± 26.5 | 0.5 ± 0.095 |
| ari-1 | -189.96 ± 15 | 0.6 ± 0.031 |
| hiw | -137.96 ± 13.9 | 0.84 ± 0.036 |
| mib1 | -174.76 ± 5 | 0.55 ± 0.027 |
| Dp | -183.78 ± 30.2 | 0.58 ± 0.03 |
| HERC2 | -189.16 ± 9.7 | 0.48 ± 0.029 |
| l(2)dtl | -151.14 ± 13.3 | 0.61 ± 0.026 |
| CG4238 | -145.46 ± 11.2 | 0.7 ± 0.03 |
| vhl | -104.45 ± 11.2 | 0.52 ± 0.087 |
| parkin | -162.5 ± 10 | 0.73 ± 0.02 |
| roc1b | -141.69 ± 6.7 | 0.81 ± 0.029 |
| Msl-2 | -102.45 ± 7.8 | 0.55 ± 0.037 |
| Sip3 | -162.2 ± 8.7 | 0.51 ± 0.078 |
| Su(dx) | -139.23 ± 13.6 | 0.73 ± 0.041 |
| Ubr1 | -148.75 ± 19.2 | 0.46 ± 0.047 |
| Topors | -80.82 ± 11.4 | 0.56 ± 0.045 |
| Morula | -140.82 ± 0.1 | 0.52 ± 0.091 |

| | | |
|---|---|---|
| Smurf1 | -136.68 ± 11 | 0.5 ± 0.006 |
| Sce line 1 | -131.91 ± 9 | 0.58 ± 0.154 |
| Sce line 2 | -130.86 ± 9.7 | 0.6 ± 0.037 |
| CG6966 | -198.42 ± 14.6 | 0.76 ± 0.071 |
| Fsn | -134.26 ± 12.2 | 0.63 ± 0.051 |
| Apc | -154.84 ± 7.7 | 0.47 ± 0.045 |
| not | -192.52 ± 13.9 | 0.69 ± 0.022 |
| godzilla | -213.81 ± 9.5 | 0.59 ± 0.042 |
| CG9003 | -127.76 ± 14 | 0.46 ± 0.02 |
| faf | -176.48 ± 15.9 | 0.76 ± 0.112 |
| CG32850 line 1 | -188.94 ± 4.3 | 0.92 ± 0.235 |
| CG32850 line 2 | -194.5 ± 7.5 | 0.37 ± 0.038 |
| Bub1 | -217.16 ± 6.2 | 0.51 ± 0.06 |
| chi line 1 | -130.49 ± 7.3 | 0.53 ± 0.098 |
| chi line 2 | -124.52 ± 23.9 | 0.57 ± 0.012 |
| CG2218 | -205.49 ± 12.6 | 0.77 ± 0.042 |
| CG15011 | -183.26 ± 19.7 | 0.58 ± 0.057 |
| CG8184 line 1 | -133.34 ± 5.2 | 0.33 ± 0.014 |
| CG8184 line 2 | -150.57 ± 22.4 | 0.5 ± 0.049 |
| CG8184 line 3 | -168.85 ± 9.9 | 0.68 ± 0.057 |
| gol | -141 ± 5.6 | 0.83 ± 0.228 |
| Shal | -166.74 ± 10.5 | 1 ± 0.359 |
| CG15800 | -184.25 ± 10.5 | 0.57 ± 0.057 |
| Nedd8 | -163.49 ± 20.8 | 0.86 ± 0.06 |
| CG42797 line 1 | -127.31 ± 9.3 | 0.61 ± 0.071 |
| CG42797 line 2 | -150.41 ± 12.8 | 0.81 ± 0.059 |
| lt | -168.17 ± 9.4 | 0.68 ± 0.032 |
| CG10440 | -132.5 ± 42.5 | 0.67 ± 0.113 |
| CG3894 | -165.33 ± 7.3 | 0.69 ± 0.035 |
| CG6179 | -132.94 ± 22.6 | 0.83 ± 0.076 |
| CG10465 | -146.5 ± 38.5 | 0.62 ± 0.153 |
| Rab40 | -202.53 ± 23.1 | 0.77 ± 0.04 |
| CG14647 | -174.53 ± 10 | 0.92 ± 0.063 |
| snky | -155.08 ± 14 | 0.84 ± 0.02 |
| mib2 | -197.39 ± 20.5 | 0.69 ± 0.06 |
| stc line 1 | -174.01 ± 21.6 | 0.62 ± 0.117 |
| stc line 2 | -180.43 ± 11.4 | 0.55 ± 0.083 |
| CG13025 | -132.89 ± 38.6 | 0.55 ± 0.089 |
| CG6752 | -196.84 ± 40.7 | 0.44 ± 0.076 |
| skpE | -147.51 ± 18.7 | 0.61 ± 0.127 |
| l(3)73Ah | -165.68 ± 8.1 | 0.46 ± 0.019 |
| CG11658 | -151.26 ± 2.2 | 0.72 ± 0.119 |
| Shaw | -177.16 ± 4.2 | 0.52 ± 0.011 |

| | | |
|---|---|---|
| CG1826 | -181.6 ± 31.7 | 0.53 ± 0.113 |
| CG5961 | -136.42 ± 20.1 | 0.79 ± 0.067 |
| Prp19 | -131.84 ± 10.3 | 0.91 ± 0.1 |
| Kdm2 | -130.92 ± 8.7 | 0.68 ± 0.031 |
| Inc | -136.65 ± 19.9 | 0.87 ± 0.037 |
| dor | -141.46 ± 5.7 | 0.83 ± 0.161 |
| Pic | -161.51 ± 9.3 | 0.52 ± 0.04 |
| TSG101 | -135.82 ± 16.5 | 0.57 ± 0.083 |
| Rbcn-3A | -181.02 ± 7.4 | 0.66 ± 0.059 |
| Poe | -157.02 ± 12.4 | 0.57 ± 0.051 |
| mat1 | -163.27 ± 22.1 | 0.85 ± 0.07 |
| Smt3 | -112.19 ± 9.1 | 0.65 ± 0.176 |
| SkpC | -144.66 ± 29.1 | 0.68 ± 0.04 |
| Mdlc | -145.75 ± 10.6 | 0.74 ± 0.118 |
| SkpA | -126.94 ± 31.4 | 0.6 ± 0.08 |
| Pnut | -184.72 ± 3.7 | 0.68 ± 0.091 |
| brat | -118.48 ± 9.3 | 0.73 ± 0.069 |
| slmb | -56.08 ± 9.5 | 0.67 ± 0.059 |
| CG17754 | -180.61 ± 8.1 | 0.75 ± 0.035 |
| Cpsf160 | -179.8 ± 19 | 0.61 ± 0.055 |
| CG9467 | -129.8 ± 2.5 | 0.64 ± 0.117 |
| CSN3 | -189.18 ± 30.8 | 0.64 ± 0.091 |
| Trc8 | -172 ± 14.2 | 0.76 ± 0.029 |
| CSN7 | -166.76 ± 10.3 | 0.53 ± 0.032 |
| CG2247 | -191.12 ± 1.8 | 0.56 ± 0.08 |
| gus | -139.8 ± 10.4 | 0.77 ± 0.068 |
| CG16952 | -227.57 ± 5.9 | 0.66 ± 0.066 |
| bon | -156.71 ± 22.6 | 0.68 ± 0.02 |
| Lpt | -199 ± 10.6 | 0.83 ± 0.072 |
| Cbl | -156.54 ± 17.2 | 0.67 ± 0.035 |
| Rbpn-5 | -198.15 ± 18.2 | 0.86 ± 0.075 |
| atl | -190.81 ± 11.1 | 0.91 ± 0.245 |
| CG4813 | -46.42 ± 14.9 | 0.88 ± 0.191 |
| CSN8 | -185.22 ± 12 | 0.9 ± 0.018 |
| alien | -174.67 ± 3.9 | 0.76 ± 0.031 |
| CSN5 | -189.7 ± 23.8 | 0.8 ± 0.113 |
| CG5382 | -126.97 ± 17.6 | 0.41 ± 0.03 |
| kel | -216.42 ± 21.9 | 0.66 ± 0.059 |
| CG11414 | -175.62 ± 23.7 | 0.67 ± 0.145 |
| CG7837 | -182.64 ± 8.6 | 0.65 ± 0.053 |
| CG11534 | -139.96 ± 6.5 | 0.84 ± 0.082 |
| mura | -110.27 ± 9.9 | 0.84 ± 0.082 |
| Fbl6 | -125.1 ± 10.7 | 0.62 ± 0.032 |

| | | |
|---|---|---|
| Mi-2 | -118.11 ± 3.7 | 0.59 ± 0.021 |
| Traf4 | -174.64 ± 15.8 | 0.62 ± 0.014 |
| Socs44A | -182.03 ± 27.1 | 0.6 ± 0.061 |
| Cnot4 | -186.61 ± 31 | 0.65 ± 0.05 |
| unk | -164.26 ± 5.3 | 0.69 ± 0.021 |
| RhoBTB | -144.82 ± 6 | 0.77 ± 0.129 |
| trx | -189.48 ± 46.3 | 0.36 ± 0.072 |
| CG1909 | -189.29 ± 17.6 | 0.72 ± 0.012 |
| d4 | -182.23 ± 0.6 | 0.62 ± 0.08 |
| CG5555 | -132.24 ± 7.5 | 0.59 ± 0.044 |
| snama | -144.77 ± 13.5 | 0.78 ± 0.082 |
| ph-p | -197.24 ± 15.3 | 0.69 ± 0.002 |
| CG32369 | -134.32 ± 7.2 | 0.57 ± 0.092 |
| Mkrn1 | -192.22 ± 5 | 0.62 ± 0.071 |
| CSN6 | -137.74 ± 7.4 | 0.6 ± 0.069 |
| dgrn | -137.34 ± 27.6 | 0.47 ± 0.091 |
| mri | -170.71 ± 13.6 | 0.82 ± 0.04 |
| pie | -230 ± 7.5 | 0.66 ± 0.089 |
| CG33144 | -131.49 ± 11.7 | 0.73 ± 0.016 |
| NSD | -130.1 ± 19.7 | 0.59 ± 0.037 |
| POSH | -111.74 ± 26.9 | 0.58 ± 0.051 |
| CG42726 | -163.23 ± 28.7 | 0.95 ± 0.069 |
| Phf7 | -136.66 ± 13 | 0.64 ± 0.031 |
| CG13442 | -138.83 ± 23.8 | 0.77 ± 0.03 |
| fancl | -113.56 ± 18.6 | 0.74 ± 0.064 |
| lmga | -150.22 ± 12.2 | 0.6 ± 0.035 |
| Ube4B | -116.11 ± 12.4 | 0.71 ± 0.054 |
| CG6923 | -95.55 ± 7.7 | 0.59 ± 0.166 |
| CG33552 | -220.47 ± 17.1 | 0.56 ± 0.075 |
| mei-P26 | -106.44 ± 8.8 | 0.51 ± 0.113 |
| lmgb | -146.68 ± 12.7 | 0.65 ± 0.04 |
| lds | -106.47 ± 4.8 | 0.81 ± 0.071 |
| Socs36E | -101.3 ± 11.9 | 0.63 ± 0.11 |
| pex10 | -137.24 ± 14.6 | 0.61 ± 0.045 |
| CG17019 | -170.18 ± 31.8 | 0.61 ± 0.045 |
| XNP | -124.12 ± 18.1 | 0.6 ± 0.088 |
| CG5071 | -211.15 ± 5.4 | 0.51 ± 0.05 |
| CG9941 | -102.55 ± 12.6 | 0.68 ± 0.021 |
| CG3711 | -119.86 ± 2.9 | 0.57 ± 0.08 |
| CSN4 | -132.68 ± 7.7 | 0.82 ± 0 |
| tth | -149.74 ± 19.4 | 0.52 ± 0.048 |
| EloA | -120.77 ± 8.1 | 0.71 ± 0.01 |
| rols | -137.59 ± 6.3 | 0.57 ± 0.055 |

| | | |
|---|---|---|
| Rabex-5 | -164.93 ± 2.7 | 0.76 ± 0.079 |
| CG10916 | -160.58 ± 23.2 | 0.73 ± 0.127 |
| ktl | -103.35 ± 14.9 | 0.72 ± 0.097 |
| CG3909 | -158.09 ± 23.2 | 0.79 ± 0.172 |
| qin | -147.59 ± 14.9 | 0.73 ± 0.063 |
| CG8786 | -138.83 ± 10 | 0.61 ± 0.17 |
| Slip1 | -210.04 ± 33.9 | 0.57 ± 0.059 |
| fbxl4 line 1 | -124.2 ± 9.8 | 0.53 ± 0.013 |
| fbxl4 line 2 | -134.42 ± 15 | 0.54 ± 0.039 |
| fbxl4 line 3 | -142.6 ± 6.6 | 0.38 ± 0.047 |
| Scrapper line 1 | -153.66 ± 12.1 | 0.43 ± 0.047 |
| Scrapper line 2 | -184.32 ± 18.9 | 0.6 ± 0.022 |

**Table S1. Summary of electrophysiology data for the genetic screen.**
Data are mean ± s.e.m. *UAS-RNAis* were driven in neurons by *elav$^{c155}$-Gal4*, and *elav$^{c155}$-Gal4*/Y served as the control. We tested 157 putative E3 ligase encoding genes and 11 E3-associated genes, using 180 lines (*UAS-RNAi* or mutants; some genes were targeted with multiple lines; mean n=4, range 3-12 per line). Control data was continuously collected throughout the genetic screen. See Methods for further details.
